# IsoPairFinder: A tool for biochemical pathway discovery using stable isotope tracing metabolomics

**DOI:** 10.1101/2025.08.18.670916

**Authors:** Zhiwei Zhou, Yuanyuan Liu, Mingxun Wang, Dylan Dodd

## Abstract

**Summary:** The functional annotation of microbial genes lags far behind genome sequencing, leaving critical gaps in our knowledge of metabolic pathways. While integrating genetic manipulation with stable isotope tracing (SIT) metabolomics holds promise for pathway discovery, existing tools lack specialized capabilities for gene perturbation experiments. To address this need, we developed IsoPairFinder, a computational tool that identifies pathway intermediates by analyzing paired unlabeled (^12^C) and isotope-labeled (^13^C) metabolomics data from gene-edited microbes. By prioritizing substrate-specific feature pairs, IsoPairFinder efficiently prioritizes biologically relevant intermediates. Implemented as an open-source R package and integrated into the GNPS2 ecosystem, IsoPairFinder provides an accessible platform for the research community to accelerate novel pathway discovery and validation.

**Availability and implementation:** IsoPairFinder is freely available as an open-source R package (https://github.com/DoddLab/IsoPairFinder) and as a web-based workflow in the GNPS2 ecosystem (https://gnps2.org/workflowinput?workflowname=isopairfinder_nextflow_workflow). Comprehensive tutorials are provided at https://doddlab.github.io/IsoPairFinder_Tutorials. These implementations support both command-line and graphical analysis workflows.

**Supplementary information:** Supplementary information is available at *Bioinformatics* online.

## 1. Introduction

Biochemical pathways are the foundation of cellular life, yet a vast number of microbial metabolic genes remain functionally uncharacterized, representing a largely unexplored frontier in biology (Pavlopoulos *et al*., 2023). This knowledge gap limits our ability to harness microbial metabolism for health, biotechnology, and sustainability. Advances in genetic tools, including CRISPR-based editing and transposon mutagenesis (Tripathi *et al*., 2024; Liu *et al*., 2020), now allow systematic disruption of microbial genes. However, connecting these genetic changes to metabolic outcomes remains challenging. Stable isotope tracing (SIT) metabolomics has the potential to address this by tracking labeled atoms through pathways, revealing intermediates that accumulate upon gene disruption. Despite its potential, existing computational tools for SIT data analysis are not optimized for gene manipulation studies (Huang *et al*., 2014; Capellades *et al*., 2016; Bueschl *et al*., 2017; Dong *et al*., 2019), leaving a need for methods that integrate genetic and metabolic insights.

To address this, we developed IsoPairFinder, a computational tool that identifies pathway intermediates by analyzing paired unlabeled (^12^C) and isotope-labeled (^13^C) metabolomics data from gene-edited microbes. By leveraging predictable mass shifts from isotopic tracers, IsoPairFinder aims to reduce false positive mass spectrometry features and prioritizes high-confidence feature pairs reflecting metabolic pathway intermediates directly affected by genetic perturbations. This approach also overcomes limitations of untargeted metabolomics in genetic disruption experiments where global metabolic changes can obscure pathway-specific signals. Additionally, IsoPairFinder provides structural clues to annotate novel intermediates, accelerating pathway discovery.

We demonstrate IsoPairFinder’s utility in a case study of gut microbial metabolism, where it identifies intermediates in genetically perturbed pathways. By integrating our tool into the GNPS ecosystem, we ensure broad accessibility to the metabolomics community. IsoPairFinder advances microbial biochemistry research and opens new avenues for engineering metabolic pathways and deciphering host-microbe interactions.

## 2. Implementation and Features

The IsoPairFinder approach is written in R and can be easily combined with available workflows. The command-line R package is freely available for direct download and installation for local applications (https://github.com/DoddLab/IsoPairFinder). To enable broader accessibility, we have also implemented IsoPairFinder within the GNPS2 ecosystem (https://gnps2.org/workflowinput?workflowname=isopairfinder_nextflow_workflow), providing a user-friendly interface for users with limited programming backgrounds.

### 2.1 SIT metabolomics with gene manipulation

IsoPairFinder identifies biochemical pathway intermediates by analyzing SIT metabolomics data from genetically perturbed systems (**Figure 1a**). The experimental design typically involves four groups: (i) wild-type (WT) microbes grown with unlabeled (^12^C) substrate, (ii) mutant microbes grown with unlabeled (^12^C) substrate, (iii) WT microbes grown with labeled (^13^C) substrate, and (iv) mutant microbes grown with labeled (^13^C) substrate. When a genetic block disrupts a pathway, two key signatures emerge in the data: (1) elevated levels of pathway intermediates in mutants compared to WT, and (2) retention-time matched isotopologue pairs (^12^C/^13^C) in mutant samples. IsoPairFinder leverages these signatures to efficiently filter LC-MS/MS (MS1 and MS2) data and pinpoint metabolic pathway intermediates.

**Figure 1.**
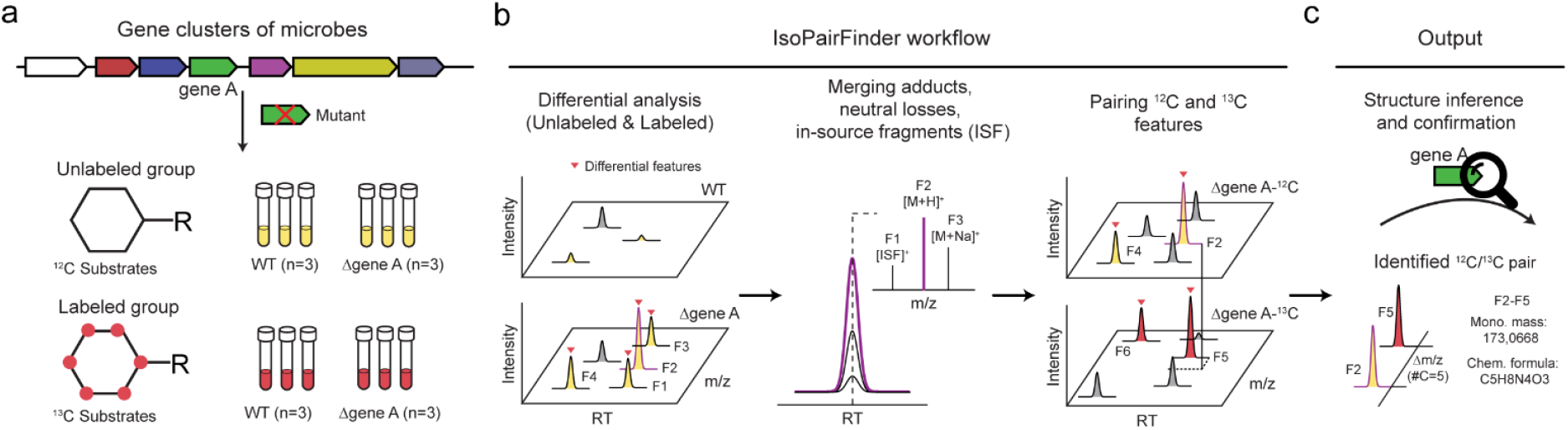
Overview of IsoPairFinder. (**a**) Schematic illustration of SIT experiments for identification of metabolic pathway intermediates; (**b**) Data processing steps of IsoPairFinder; (**c**) Output of IsoPairFinder. The abbreviation F1 represents feature 1, and ISF represents the in-source fragment.

### 2.2 Workflows of IsoPairFinder

In general, IsoPairFinder identifies the substrate feature pairs in the follow five steps (**Figure 1b-c**, and **Supplementary Figure 1**). The default parameters indicated below are adjustable according to the user’s study design.

#### 1) Data import

IsoPairFinder requires four different types of input files: feature tables, sample information table, raw MS1 data files, and raw MS2 (MS/MS) data files. The feature tables (comma-separated CSV files; unlabeled WT vs. unlabeled mutant and labeled WT vs. labeled mutant) contain differential feature results exported from common peak picking tools (e.g., XCMS (Smith *et al*., 2006), MS-DIAL (Tsugawa *et al*., 2020), and MZmine (Schmid *et al*., 2023) etc.) with minor formatting modifications. The sample information table (CSV file) defines the tracers and group information. The raw MS1 (mzML/mzXML) and MS2 (mzML/mzXML/MGF) data files are used to merge adducts, neutral loss, and in-source fragments. The detailed format requirements are provided in the tutorials.

#### 2) Differential analysis

IsoPairFinder begins by performing differential analysis to identify potential pathway intermediates affected by genetic perturbation. IsoPairFinder analyzes both unlabeled and labeled sample groups separately based on experimental design metadata provided by the user. For each group, candidate features are selected based on statistical significance (P-value < 0.05) and magnitude of change (fold-change > 10). The analysis employs Student’s t-test and false discovery rate adjustments (Benjamini & Hochberg) to compare feature intensities between WT and mutant samples. This initial screening step helps focus subsequent analyses on the most biologically relevant changes resulting from genetic perturbations.

#### 3) Merging adducts, neutral losses, and in-source fragments

Electrospray ionization produces a large number of ions, including adducts, neutral losses, and in-source fragments (ISF) that create redundancies and false positives in feature pairing. IsoPairFinder addresses this by systematically identifying and merging these ions using established methods (Zhou *et al*., 2022). The process begins by selecting differential features (WT vs. mutant) from the unlabeled group as base features. For each base feature (limited to protonated or deprotonated ions), the tool retrieves related features with a ±3 s retention time window. These feature groups are then analyzed to identify specific ion types: 41 possible adducts and 58 neutral loss types are detected based on m/z tolerances (±25 ppm), while ISFs are identified through MS/MS spectral matching and peak shape correlation analysis (Guo *et al*., 2021). After recognition, all redundant features are merged and removed from the intermediate feature list prior to pairing analysis.

#### 4) Pairing ^12^C and ^13^C features

IsoPairFinder completes the analysis by pairing differential features between unlabeled (^12^C) and labeled (^13^C) groups to identify pathway intermediate feature pairs. The tool first predicts the chemical formula of each unlabeled feature using its exact mass and the seven golden rules (Kind and Fiehn, 2007), considering only common biological elements (CHNOPS). This predicted chemical formula then determines the possible labeled atoms and calculates the theoretical m/z for corresponding ^13^C-labeled features. Finally, the algorithm matches unlabeled and labeled features by comparing their observed m/z values and retention times to these theoretical predictions, using tolerances of 10 ppm for m/z and 3 seconds for retention time. This systematic approach ensures accurate identification of biologically relevant isotope pairs.

#### 5) Output of IsoPairFinder

IsoPairFinder generates comprehensive results to support downstream analysis. The tool exports identified feature pairs with detailed information, including m/z values, retention times, mass differences, potential labeled atoms, and predicted formula. Additionally, it provides the differential analysis results along with recognized adducts, neutral loss, and in-source fragments for reference. To facilitate visual verification, IsoPairFinder creates mirror extracted ion chromatograms of the feature pairs, enabling users to efficiently review and select the most likely biologically relevant matches. These outputs serve as valuable resources for both structural inference of metabolites and experimental validation studies.

### 2.3 Comparison of SIT metabolomics tools

IsoPairFinder offers several unique advantages compared to existing SIT metabolomics tools (Huang *et al*., 2014; Capellades *et al*., 2016; Bueschl *et al*., 2017; Dong *et al*., 2019) (**Supplementary Table 1)**. First, it is specifically designed to analyze metabolomics data from experiments combining genetic manipulation with SIT, enabling identification of pathway intermediate feature pairs. Second, the tool incorporates multiple functions to enhance result reliability, including automated merging of adducts, neutral losses, and in-source fragments, along with chemical formula prediction. Third, IsoPairFinder provides flexibility by supporting output from various raw data processing platforms such as XCMS, MS-DIAL, and MZmine. This compatibility allows researchers to use their preferred data processing tools while avoiding the need for extensive parameter optimization.

## 3. Case study

We evaluated IsoPairFinder’s performance using data from our recent investigation of uric acid (UA) catabolism in gut bacteria (Liu *et al*., 2023, Liu *et al*., 2025). This study aimed to characterize the metabolic pathway by which gut bacteria degrade UA into short-chain fatty acids, a process that compensates for the loss of uricase in hominids (Liu *et al*., 2025). Using CRISPR and ClosTron, we generated seven single-gene mutant strains of *Clostridium sporogenes* (**Supplementary Figure 2a**). WT and mutant strains were cultured with either unlabeled UA or [^13^C_5_]-labeled UA for 48 hours, producing four experimental groups for analysis.

LC-MS/MS analysis with HILIC columns identified the potential pathway intermediates (**Supplementary Figure 2b**). After processing raw data with MS-DIAL, we applied IsoPairFinder to identify reliable feature pairs. The tool effectively filtered 50-99% of theoretical feature pairs, significantly reducing false positives (**Supplementary Figure 3** and **Supplementary Data 1**). For instance, in *ygeW* mutant strains, IsoPairFinder narrowed 725 theoretical feature pairs down to four high-confidence candidates. We selected the most abundant pair (191.0747@5.417_196.0914@5.414) for validation, whose mass difference indicated incorporation of five carbon atoms. Chemical formula prediction (C5H10N4O4) and functional analysis suggested the structure as 2,3-diureidopropanoic acid.

Through this approach, we successfully determined 4 of 6 substrate structures using chemical standards and reconstructed the UA catabolism pathway in vitro (Liu *et al*., 2025). These results demonstrate IsoPairFinder’s effectiveness in identifying biologically relevant intermediate feature pairs for biochemical pathway discovery using SIT metabolomics.

## 4. Conclusion and Discussion

IsoPairFinder is a freely available computational tool designed to identify the pathway intermediate feature pairs in SIT metabolomics studies. By integrating genetic manipulation techniques, it enables researchers to efficiently screen candidate features and prioritize biologically relevant intermediates for validation. In our case study, we demonstrated its utility for characterizing a novel metabolic pathway in gut bacteria. However, IsoPairFinder is equally applicable to other biological systems with established genetic tools, including mammalian cells and model microorganisms, where it could facilitate discovery of novel natural products, metabolic pathways, and biological mechanisms.

While IsoPairFinder effectively identifies feature pairs at the MS1 level, structural elucidation requires complementary approaches. Future integration with MS/MS analysis tools like SIRIUS (Dührkop *et al*., 2019) and MS/MS analog searching could enhance IsoPairFinder’s capabilities for comprehensive metabolite characterization. For optimal performance, we recommend using ^13^C isotope labeling in the IsoPairFinder workflow to minimize chromatographic retention time shifts of pathway intermediates.

## Acknowledgements

We thank all lab members for their discussions to improve IsoPairFinder.

## References

Bueschl, C. et al. (2017) MetExtract II: A Software Suite for Stable Isotope-Assisted Untargeted Metabolomics. Anal. Chem., 89, 9518–9526.

Capellades, J. et al. (2016) geoRge: A Computational Tool To Detect the Presence of Stable Isotope Labeling in LC/MS-Based Untargeted Metabolomics. Anal. Chem., 88, 621–628.

Dong, Y. et al. (2019) Miso: an R package for multiple isotope labeling assisted metabolomics data analysis. Bioinformatics, 35, 3524–3526.

Dührkop, K. et al. (2019) SIRIUS 4: a rapid tool for turning tandem mass spectra into metabolite structure information. Nat Methods, 16, 299–302.

Guo, J. et al. (2021) ISFrag: De Novo Recognition of In-Source Fragments for Liquid Chromatography–Mass Spectrometry Data. Anal. Chem., 93, 10243–10250.

Huang, X. et al. (2014) X13CMS: Global Tracking of Isotopic Labels in Untargeted Metabolomics. Anal. Chem., 86, 1632–1639.

Kind, T. and Fiehn, O. (2007) Seven Golden Rules for heuristic filtering of molecular formulas obtained by accurate mass spectrometry. BMC Bioinformatics, 8.

Liu, Y. et al. (2023) A widely distributed gene cluster compensates for uricase loss in hominids. Cell, 186, 3400-3413.e20.

Liu, Y. et al. (2025) Gut bacteria degrade purines via the 2,8-dioxopurine pathway. Nat Microbiol.

Liu, Z. et al. (2020) Application of different types of CRISPR/Cas-based systems in bacteria. Microb Cell Fact, 19.

Pavlopoulos, G.A. et al. (2023) Unraveling the functional dark matter through global metagenomics. Nature, 622, 594–602.

Schmid, R. et al. (2023) Integrative analysis of multimodal mass spectrometry data in MZmine 3. Nat Biotechnol, 41, 447–449.

Smith, C.A. et al. (2006) XCMS: Processing Mass Spectrometry Data for Metabolite Profiling Using Nonlinear Peak Alignment, Matching, and Identification. Anal. Chem., 78, 779– 787.

Tripathi, S. et al. (2024) Randomly barcoded transposon mutant libraries for gut commensals I: Strategies for efficient library construction. Cell Reports, 43, 113517.

Tsugawa, H. et al. (2020) A lipidome atlas in MS-DIAL 4. Nat Biotechnol, 38, 1159–1163.

Zhou, Z. et al. (2022) Metabolite annotation from knowns to unknowns through knowledge-guided multi-layer metabolic networking. Nat Commun, 13, 6656.

