## Supplementary Information for "IsoPairFinder: A tool for biochemical pathway discovery using stable isotope tracing metabolomics"

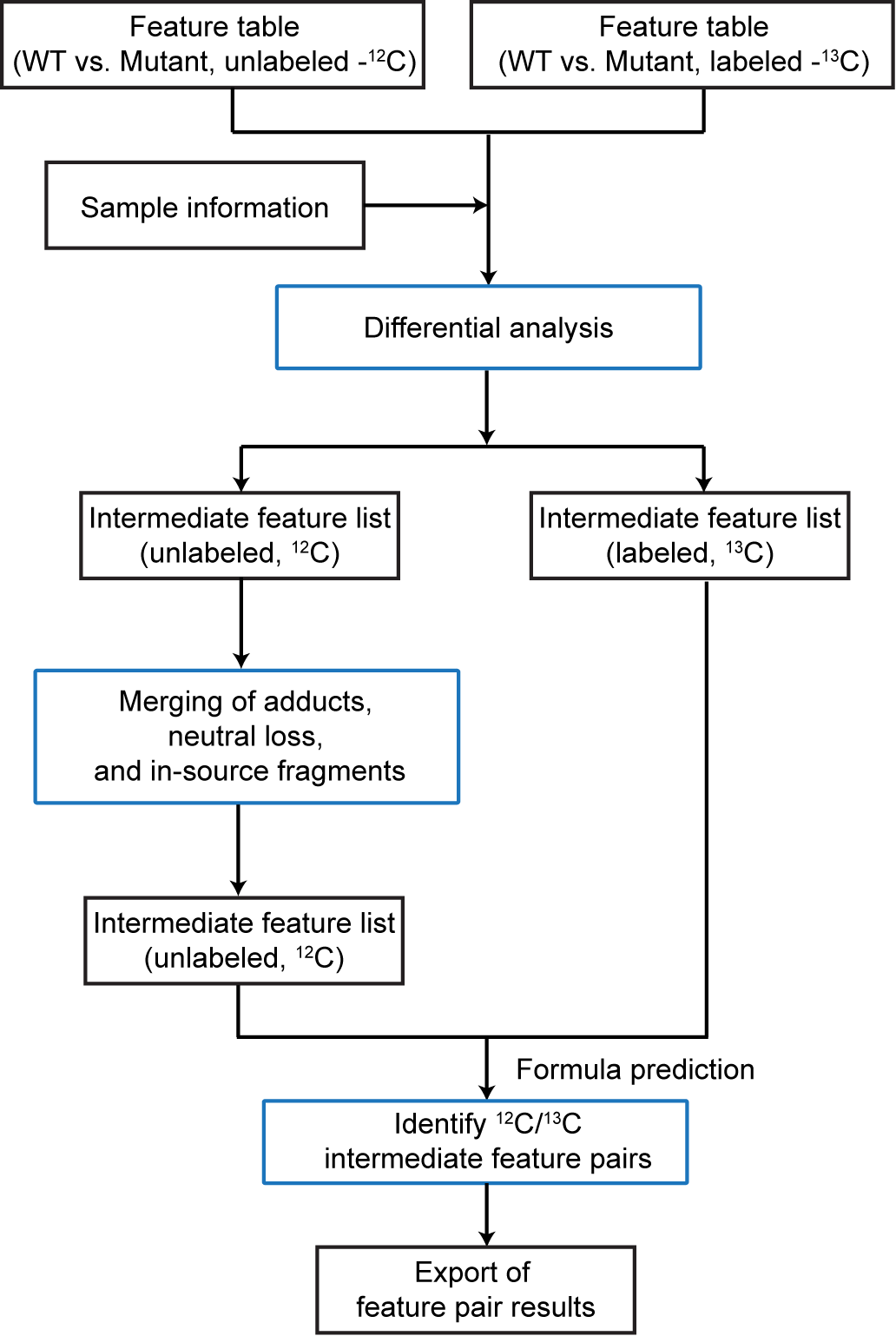

**Supplementary Figure 1**: Flow chart of the IsoPairFinder data processing.

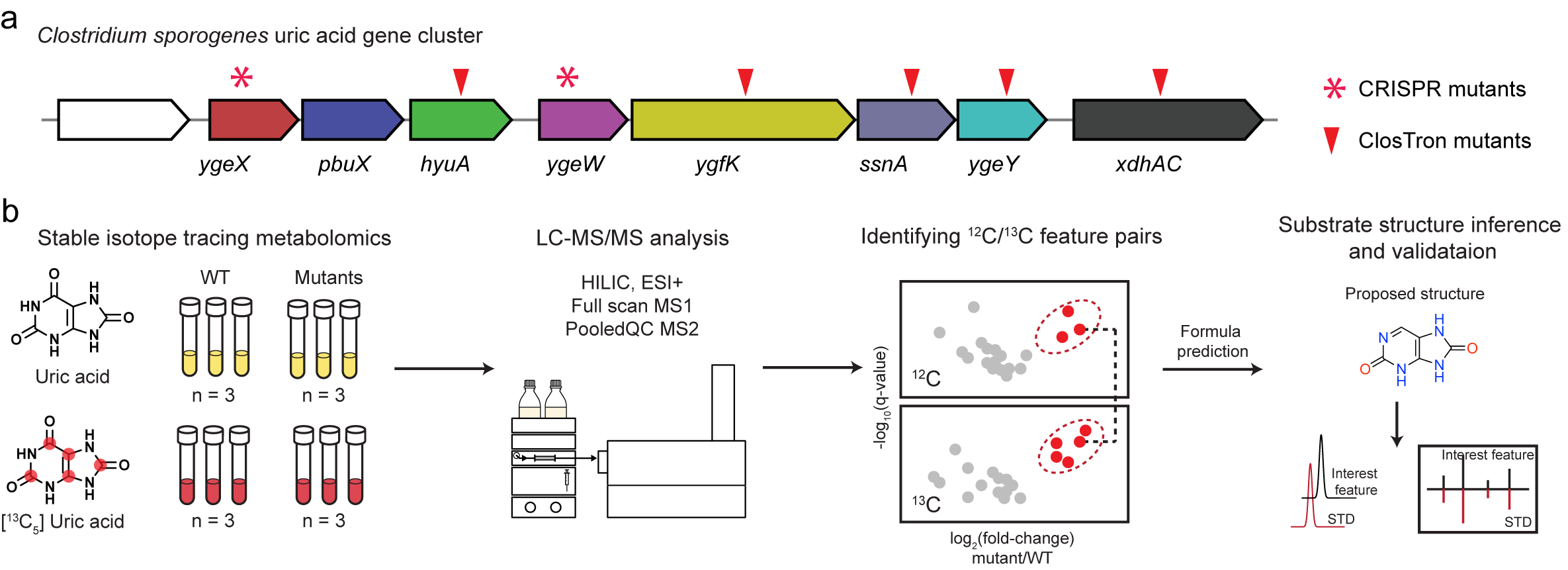

**Supplementary Figure 2**: Case study using IsoPairFinder to uncover a previously unknown pathway for uric acid catabolism. (**a**) CRISPR and ClosTron mutants constructed in a uric acid-inducible gene cluster in *Clostridium sporogenes*; (**b**) Schematic illustration of how IsoPairFinder is used to investigate novel intermediates in uric acid catabolism.

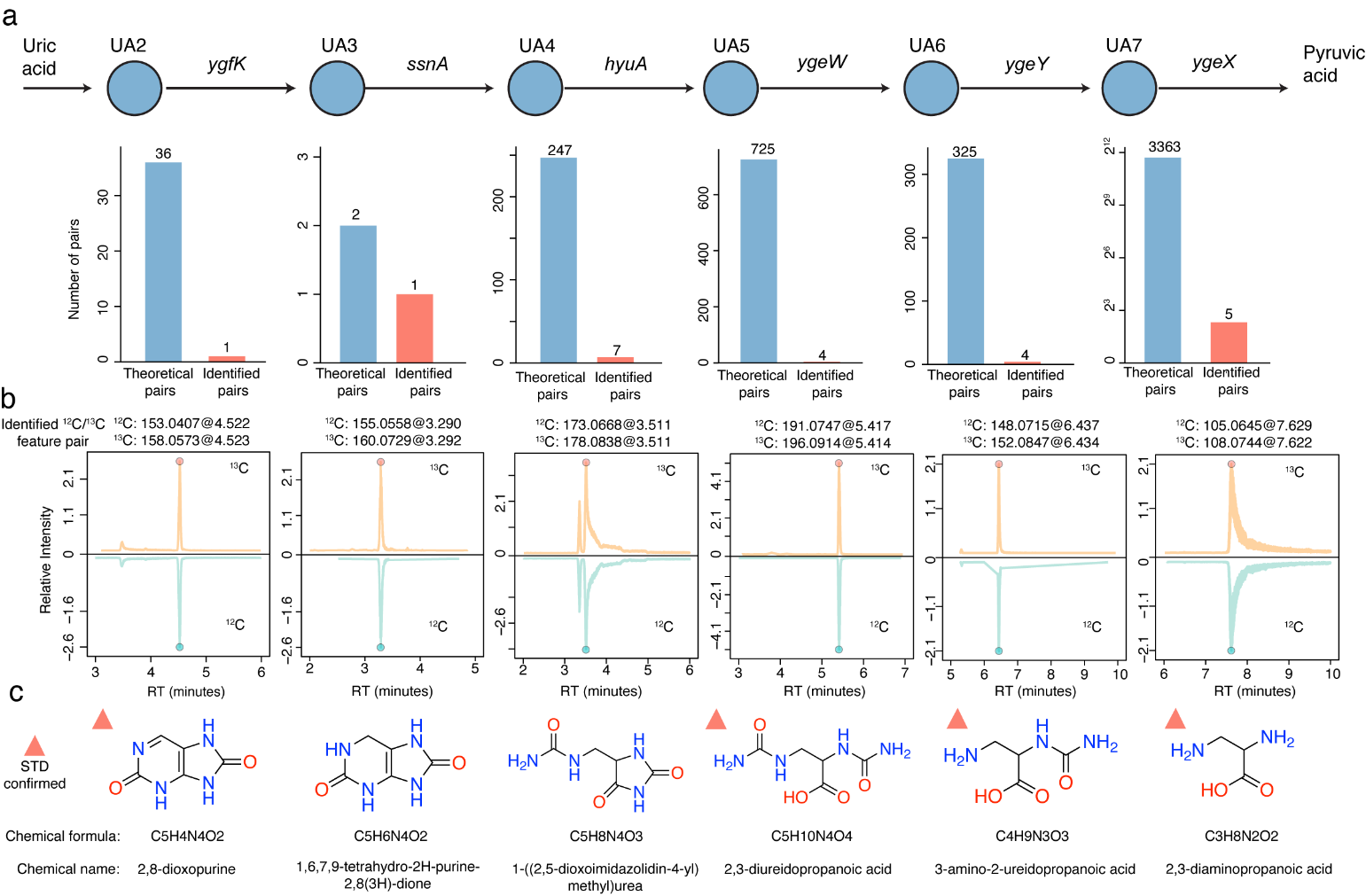

**Supplementary Figure 3**: Identification of ^12^C/^13^C feature pairs in uric acid catabolism. (**a**) IsoPairFinder reduces the number of feature pairs identified in metabolomics data from wild-type and mutants during uric acid degradation; (**b**) Extracted ion chromatograms (EICs) for most abundant ^12^C/^13^C feature pairs identified by IsoPairFinder; (**c**) Proposed and confirmed structures of metabolic intermediates in the uric acid degradation pathway.

**Supplementary Table 1.** Comparison of available tools for stable isotope tracing metabolomics data processing

|  | IsoPairFinder | X13CMS  (Huang *et al.*, 2014) | geoRge (Capellades *et al.*, 2016) | MetExtract (Bueschl *et al.*, 2017) | Miso  (Dong *et al.*, 2019) |
| --- | --- | --- | --- | --- | --- |
| Objective | Identify pathway intermediates from gene mutants | Track isotopologue distribution changes | Track isotopologue distribution changes | Detect isotopically labeled molecules | Detect isotopically labeled molecules |
| Designed to integrate genetic perturbations | Yes | No | No | No | No |
| Output | Feature pairs | Isotopologue groups | Isotopologue groups | Feature pairs | Feature pairs |
| Merging adducts, neutral losses, in-source fragments | +++ | No | No | + | No |
| Formula prediction | Yes | No | No | No | No |
| Compatibility | +++  (XCMS, MS-DIAL, MZmine) | +  (XCMS) | +  (XCMS) | -  (Independent) | +  (XCMS) |
| Graphical User Interface (GUI) | Yes | No | No | Yes | No |

References

Bueschl,C. *et al.* (2017) MetExtract II: A Software Suite for Stable Isotope-Assisted Untargeted Metabolomics. *Anal. Chem.*, **89**, 9518–9526.

Capellades,J. *et al.* (2016) geoRge: A Computational Tool To Detect the Presence of Stable Isotope Labeling in LC/MS-Based Untargeted Metabolomics. *Anal. Chem.*, **88**, 621–628.

Dong,Y. *et al.* (2019) Miso: an R package for multiple isotope labeling assisted metabolomics data analysis. *Bioinformatics*, **35**, 3524–3526.

Huang,X. *et al.* (2014) X^13^CMS: Global Tracking of Isotopic Labels in Untargeted Metabolomics. *Anal. Chem.*, **86**, 1632–1639.
